# Lonely in the dark: trauma memory and sex-specific dysregulation of amygdala reactivity to fear signals

**DOI:** 10.1101/2021.11.16.468598

**Authors:** Mitjan Morr, Jeanine Noell, Daphne Sassin, Jule Daniels, Alexandra Philipsen, Benjamin Becker, Birgit Stoffel-Wagner, René Hurlemann, Dirk Scheele

## Abstract

Loneliness exacerbates psychological distress and increases the risk of psychopathology after trauma exposure. The prevalence of trauma-associated disorders varies substantially between sexes, and accumulating evidence indicates sex-specific effects of loneliness. However, it is still unclear whether a lack of social connectedness affects trauma-induced intrusions and the neural processing of fear signals. Moreover, it is uncertain, whether loneliness plays a different role in women and men. We used a prestratification strategy and recruited n=47 (n=20 women) healthy individuals with high loneliness and n=35 controls (n=18 women). Participants were exposed to an experimental trauma and evoked intrusive thoughts in daily life were monitored for three consecutive days. Functional magnetic resonance imaging was used to assess neural habituation to fearful faces and fear learning (conditioning and extinction) prior to trauma exposure. The total number of intrusions and amygdala reactivity in neural fear processing served as the primary study outcomes. Our results revealed a significant interaction between loneliness and sex such that loneliness was associated with more intrusions in men, but not in women. A similar pattern emerged at the neural level, with both reduced amygdala habituation to repeated fearful faces and amygdala hyperreactivity during the conditioning of fear signals in lonely men, but not in women. Our findings indicate that loneliness may confer vulnerability to intrusive memories after trauma exposure in healthy men and that this phenotype relates to altered limbic processing of fear signals. Collectively, interventions targeting social connectedness may mitigate the sequelae of traumatic experiences.

## Introduction

Loneliness, defined as the discrepancy between desired and actual social connectedness [1], is a growing problem in modern societies and has been further exacerbated by the COVID-19 pandemic [2–5]. Loneliness can be considered as the social equivalent to hunger or pain to meet social needs and has been associated with increased mortality, resembling risk factors like obesity or smoking [6, 7]. Furthermore, loneliness is closely linked with various psychiatric disorders such as substance abuse, depression and anxiety disorders [8, 9]. Importantly, loneliness also constitutes a risk factor for developing posttraumatic stress disorder (PTSD) following a traumatic experience [10, 11]. In fact, loneliness predicts future PTSD and is predicted by past PTSD symptoms, indicating a bidirectional relationship between PTSD and social connectedness [12, 13].

PTSD is a debilitating and frequently chronic condition characterized by intrusive thoughts about the traumatic experience as a key symptom [14–16]. Intrusions are defined as involuntarily spontaneous memories of the distressing incident, mainly experienced as visual forms of mental imagery [17–19]. The lifetime prevalence of PTSD varies substantially between sexes, with women being twice as likely to develop PTSD than men [20]. Current neurocircuit models of PTSD highlight dysfunction of the amygdala-hippocampus complex as a core mechanism underlying the persistence of intrusive memories. Modern trauma-focused psychotherapies for treating intrusions and other PTSD symptoms often include an exposure-based intervention to reduce fear responses [21]. Mechanistically, this decrease in fear responses can be achieved by both fear extinction and fear habituation. The former is characterized by a progressive decrement of a conditioned fear response (CR) when a conditioned stimulus (CS) is repeatedly presented in the absence of an aversive unconditioned stimulus (UCS) with which it has previously been paired, while the latter is based on repeated exposure to the (imagined) UCS. In fact, both fear extinction and habituation recruit overlapping forebrain structures including the amygdala [22].

The experimental trauma paradigm is a widely used and reliable method to evoke intrusions by showing traumatizing film footage in a controlled laboratory setting [17, 23, 24]. On a neural level, increased reactivity in the amygdala, hippocampus, insula and anterior cingulate cortex (ACC) during trauma exposure predicts increased intrusive thoughts [25, 26]. Interestingly, neural processing during fear extinction has also been linked to intrusion frequency in an experimental trauma paradigm and reduced extinction capacity predicts PTSD development [27, 28]. Furthermore, women reported more intrusive symptoms following the trauma paradigm than men, and this sex difference was related to peritraumatic responding and slowed extinction learning in women [29]. Likewise, women showed a sustained amygdala response to negative evocative images relative to men [30].

We previously found that strong trauma disclosure reduces intrusions and alters amygdala functional connectivity following trauma exposure only in individuals with heightened concentrations of the hypothalamic peptide oxytocin [31]. Given a crucial role of oxytocin in safety learning and a reduced oxytocin reactivity to positive social interactions in people experiencing loneliness [32, 33], this raises the intriguing possibility that loneliness influences intrusions after trauma exposure by modulating self-disclosure and amygdala-related fear processing. Furthermore, a recent large-scale study indicated a higher prevalence of loneliness in men than in women [34], and a growing number of studies reported sex-specific effects of loneliness. For instance, loneliness was associated with more pronounced within-network coupling of the default network in men than in women, and brain volume effects in the limbic system were linked to the frequency and intensity of social contact in a sex-dependent manner [35, 36]. Surprisingly, however, the impact of loneliness on fear conditioning/extinction and fear habituation as well as the possible moderation by sex remain unclear. Therefore, this study aimed to examine loneliness-associated neurobiological risk factors for intrusive thoughts in an experimental prospective study design.

To this end, we recruited a prestratified sample of 82 healthy volunteers assigned to either a high-lonely and low-lonely group to test how loneliness interacts with sex to influence the neural processing of fear signals and the formation of intrusive thoughts. During functional magnetic resonance imaging (fMRI), subjects completed an emotional face-matching task to assess neural responses to fearful faces and the habituation of these responses, as well as a fear conditioning and extinction paradigm. Subsequently, we probed psychological, physiological and hormonal responses in an experimental trauma paradigm and measured the evoked intrusions during the following three days after trauma exposure. We hypothesized that lonely individuals would exhibit more pronounced responses to the experimental trauma film and experience more intrusions. Furthermore, we expected to observe loneliness-dependent hyperreactivity to fearful faces and fear-conditioned stimuli in the amygdala, as well as changes in functional connectivity in a network responsible for fear processing [37–39]. Given previous findings about sex differences in the effects of loneliness and the formation of intrusive memories, we explored sex as a moderator variable.

## Materials and methods

### Participants

The present study used a quasi-experimental design with a sample of prestratified healthy volunteers scoring high or low on the revised UCLA Loneliness Scale (UCLA LS) [40]. High scorers (high-lonely) were defined by a score above or equal to 50 (i.e., at least one standard deviation above the mean score of healthy young adults [41], which is similar to previous categorizations [42]), while low scorers (low-lonely) were defined by a score of 25 or below (i.e., at least one standard deviation below the mean). In total, 4515 participants completed the UCLA LS online questionnaire and clinical interview were conducted with 97 subjects. The final sample consisted of 82 healthy subjects (mean age ± standard deviation [SD]: 26.39 ± 5.83 years) assigned to either a high-loneliness (n = 47 [20 women]) and a low-loneliness control group (n = 35 [18 women]). All participants gave written informed consent. The study was approved by the institutional review board of the medical faculty of the University of Bonn and carried out in compliance with the latest revision of the Declaration of Helsinki.

### Experimental design

In screening sessions, medical history and psychiatric symptoms were assessed (shown in supplementary information (SI) for inclusion criteria and **Figure S1** for a design overview). The testing session consisted of an fMRI scan containing a high-resolution structural scan, a fear conditioning (COND) / extinction (EXT) paradigm [43], and a well-established emotional face-matching paradigm [44]. All magnetic resonance imaging (MRI) data were acquired using a 3T Siemens TRIO MRI system (Siemens AG, Erlangen, Germany) with a Siemens 32-channel head coil. Following fMRI acquisition, the participants completed an experimental trauma paradigm [45]. To measure trauma disclosure and intrusive thoughts, subjects completed online diaries during the following three days after trauma exposure. Saliva samples were collected before the fMRI scan as baseline measure, and before and after the experimental trauma paradigm to measure oxytocin levels. In addition, blood samples were taken before the fMRI scan to measure the levels of gonadal steroids including estradiol and testosterone, as control variables. For a detailed list of the questionnaires and neuroendocrine parameters, see SI.

### Emotional face matching task

The first fMRI paradigm consisted of an adapted version of a well-established emotional face-matching paradigm [44, 46]. Subjects had to match two simultaneously presented pictures at the bottom with a target picture presented at the top of the screen. Stimuli consisted of pictures of faces (neutral, fearful, and happy) and houses as non-social control stimuli. Stimuli were presented in three blocks for every condition (happy, fearful, and neutral faces, as well as houses), with each block consisting of five trials. Participants had to match the face identity (i.e., the emotion was consistent across all faces of a trial).

### Fear conditioning and extinction tasks

In the COND phase, subjects were shown four different pictures (two neutral faces [social stimuli] and two houses [non-social stimuli]). Within each category, one picture was designated as fear-associated CS (CS+) and the other as safety signal (CS-). Each stimulus was presented 16 times during the COND and EXT experiments. In 75% of CS+ trials, subjects received an electric impulse (the UCS) 4 s after stimulus onset. The electric impulses were delivered by a Biopac System (MP150, Biopac Systems Inc., Goleta USA). In addition, the Biopac system measured electrodermal activity (EDA) and respiration during the experiment. No electrical impulses were administered in the EXT phase. In both phases, participants had to press a button before the UCS to indicate if they believed that they would receive an electric impulse (i.e., a contingency rating was coded by +1 for an expected shock and −1 for no shock). For a detailed description of the data acquisition, preprocessing, and analyses of both tasks, see the SI.

### Experimental trauma paradigm

Participants were seated in front of a Tobii TX300 binocular eye-tracker (Tobii AB, Danderyd, Sweden) with a 23-inch display to measure pupil sizes as the outcome indicating physical arousal during the movie alongside EDA. To evoke intrusive thoughts, participants were confronted with a 24-minute-long movie clip derived showing the multiple rape of a young woman by a group of men. EDA data were measured with a Biopac MP150 system. Positive and negative affect, dissociative symptoms, valence (0 = low valence, 100 = high valence), arousal (0 = low arousal, 100 = high arousal) as well as state anxiety were measured prior and after the experimental trauma paradigm. The participants completed online intrusion diaries at home in the evening during three consecutive days following trauma exposure. The intrusion diaries included, inter alia, the number of intrusions, ratings of intrusion-induced stress, and whether the subjects discussed or wanted to discuss the trauma movie with other people. For details about data collection and preprocessing, see the SI.

### Statistical analyses

Our primary outcomes included the number of intrusions and blood oxygen level-dependent (BOLD) signal changes during fear learning and fear habituation. Other outcomes recorded were the psychological and physiological stress markers after trauma exposure and skin conductance response during fear conditioning. Mixed-design analyses of variance (ANOVAs) and Bonferroni-corrected (*p*_cor_) post hoc *t*-tests were calculated using SPSS 25 to examine changes in intrusive thoughts (sum of the three consecutive days following the trauma exposure), trauma disclosure (i.e., how long participants talked to other people and whether and how long they discussed the trauma movie with other people), group differences in psychiatric symptoms and psychological as well as physiological and hormonal responses to the trauma exposure with the between-subject factors of sex (women, men) and loneliness (high, low). Mixed-design ANOVAs for contingency ratings included the additional within-subject factors task (COND, EXT) and condition (CS+, CS-). Additional mixed-design ANOVAs for the COND/EXT paradigms included the between-subject factors of sociality (social, non-social) and time (first half, second half).

To analyze the fMRI data, we used a two-stage approach as implemented in SPM12 (Wellcome Trust Center for Neuroimaging, London, UK; http://www.fil.ion.ucl.ac.uk/spm). On the first level, data were modeled using a fixed-effects model. On the second level, the main contrasts of interest were compared between groups using a full factorial model with the two factors of loneliness and sex. Analyses were conducted using anatomically defined regions of interest (ROIs), including the amygdala, derived from the WFU PickAtlas (for further ROI results, see the SI). *p*-values < 0.05 after familywise error correction for multiple testing (*p*_FWE_) were considered significant. In addition, generalized psychophysiological interaction (gPPI) analysis was conducted to assess functional connectivity by using the CONN toolbox 18.a (www.nitrc.org/projects/conn, RRID:SCR_009550) [47]. Pearson correlations between parameter estimates of significant ROI clusters and intrusive thoughts were calculated.

## Results

### Subclinical psychiatric symptoms, loneliness, and sex differences

High-lonely subjects reported more depressive symptoms, alexithymia, childhood maltreatment, social interaction anxiety, and subjective stress compared to low-lonely participants (all *p*s < 0.02; shown in **Tab. 1**). Furthermore, high-lonely participants had smaller and less diverse social networks and received less social support (all *p*s < 0.03). In addition, across groups, women reported having more social support than men (*F*_(1,78)_ = 5.12, *p* = 0.03, η_p_^2^ = 0.06). There were no significant interactions between sex and loneliness in psychiatric symptoms and social network quality (all *p*s > 0.05). Besides the expected sex differences, we found a significant sex*loneliness interaction in estradiol levels (*F*_(1,65)_ = 7.60, *p* = 0.01, η_p_^2^ = 0.11), showing that high-lonely women exhibited higher estradiol levels than low-lonely women at the fMRI session (*t*_(16.55)_ = 2.62, *p*_cor_ = 0.04, *d* = 0.87; shown in **Tab. S1**).

**Table 1.** Baseline differences between the high-lonely and low-lonely groups

|  | High-lonely<br>(*n* = 47) | Low-lonely<br>(*n* = 35) | *t* | *p* |
| --- | --- | --- | --- | --- |
| Loneliness <sup>a)</sup> | 54.94 (4.49) | 23.80 (1.13) | 45.60 | < 0.01 |
| Depressive symptoms <sup>b)</sup> | 4.02 (3.71) | 1.83 (2.99) | 2.96 | < 0.01 |
| Social anxiety <sup>c)</sup> | 22.38 (18.05) | 12.63 (12.68) | 2.87 | 0.01 |
| Childhood maltreatment <sup>d)</sup> | 36.98 (9.84) | 30.83 (11.51) | 2.60 | 0.01 |
| Alexithymia <sup>e)</sup> | 44.06 (10.27) | 33.31 (6.48) | 5.79 | < 0.01 |
| Social support <sup>f)</sup> | 55.64 (12.18) | 66.89 (9.20) | -4.77 | < 0.01 |
| Perceived stress <sup>g)</sup> | 13.09 (6.67) | 8.09 (4.87) | 3.93 | < 0.01 |
| Trait anxiety <sup>h)</sup> | 38.79 (9.03) | 27.02 (4.93) | 7.55 | < 0.01 |
| Social network <sup>i)</sup> |  |  |  |  |
| Number of people | 15.87 (7.47) | 20.31 (7.40) | 2.67 | 0.01 |
| Roles | 4.87 (1.33) | 5.71 (1.51) | 2.68 | 0.01 |
| Networks | 1.53 (1.20) | 2.14 (1.16) | 2.36 | 0.02 |

*Notes.* Values are the mean and SD. <sup>a)</sup> Participants were prestratified and assigned to the high- or low-lonely group using the UCLA Loneliness Scale (UCLA-L). High-lonely participants had a score equal or above 50, while low-lonely participants had a score equal or below 25; <sup>b)</sup> Depressive symptoms were measured with the Beck Depression Inventory, Version II (BDI); <sup>c)</sup> Social anxiety was assessed with the Liebowitz Social Anxiety Scale (LSAS); <sup>d)</sup> Childhood traumata were measured using the Childhood Trauma Questionnaire (CTQ); <sup>e)</sup> Alexithymic symptoms were assessed by the Toronto Alexithymia Scale (TAS); <sup>f)</sup> Social Support was measured with the Social Support Questionnaire ([Fragebogen zur sozialen Unterstützung] ;F-SozU); <sup>g)</sup> Perceived stress was quantified by the Perceived Stress Scale (PSS-10); <sup>h)</sup> Trait anxiety was assessed by the State Trait Anxiety Inventory (STAI); <sup>i)</sup> Social network was characterized using the Social Network Index assessing the number of diverse social roles, networks, and the total number of people to whom the participants talk to regularly.

### Psychological and physiological reaction to the trauma video

After trauma exposure, subjects showed dissociative symptoms (mean ± SD = 1.24 ± 1.18, one-sample t-test against zero: *t*_(77)_ = 9.36, *p* < 0.01, *d* = 1.06) and reported high arousal (76.87 ± 23.53) induced by and low valence (9.35 ± 16.16) of the trauma film. Neither dissociative symptoms nor valence and arousal were affected by loneliness or sex (all *p*s > 0.05). Subjects showed a decrease in positive affect (main effect of time: *F*_(1,72)_= 67.88, *p* < 0.01, η_p_^2^ = 0.49; shown in **Fig. 1A**) and an increase in negative affect (main effect of time: *F*_(1,72)_= 139.58, *p* < 0.01, η_p_^2^ = 0.66; shown in **Fig. 1B**) independent of sex and loneliness following the trauma video. In addition, state anxiety increased significantly (main effect of time: *F*_(1,72)_ = 154.91, *p* < 0.01, η_p_^2^ = 0.68; shown in **Fig. 1C**) and we observed an interaction between loneliness and time (*F*_(1,72)_ = 4.44, *p* = 0.04, η_p_^2^ = 0.06), such that lonely individuals displayed higher baseline state anxiety ratings (*t*_(76)_ = 4.42, *p*_cor_ < 0.01, *d* = 1.02) than low-lonely individuals, but state anxiety significantly increased in both groups (high-lonely: *t*_(41)_ = 8.98, *p* < 0.01, *d* = 1.39; low-lonely: *t*_(33)_ = 7.99, *p* < 0.01, *d* = 1.37).

**Fig. 1.**
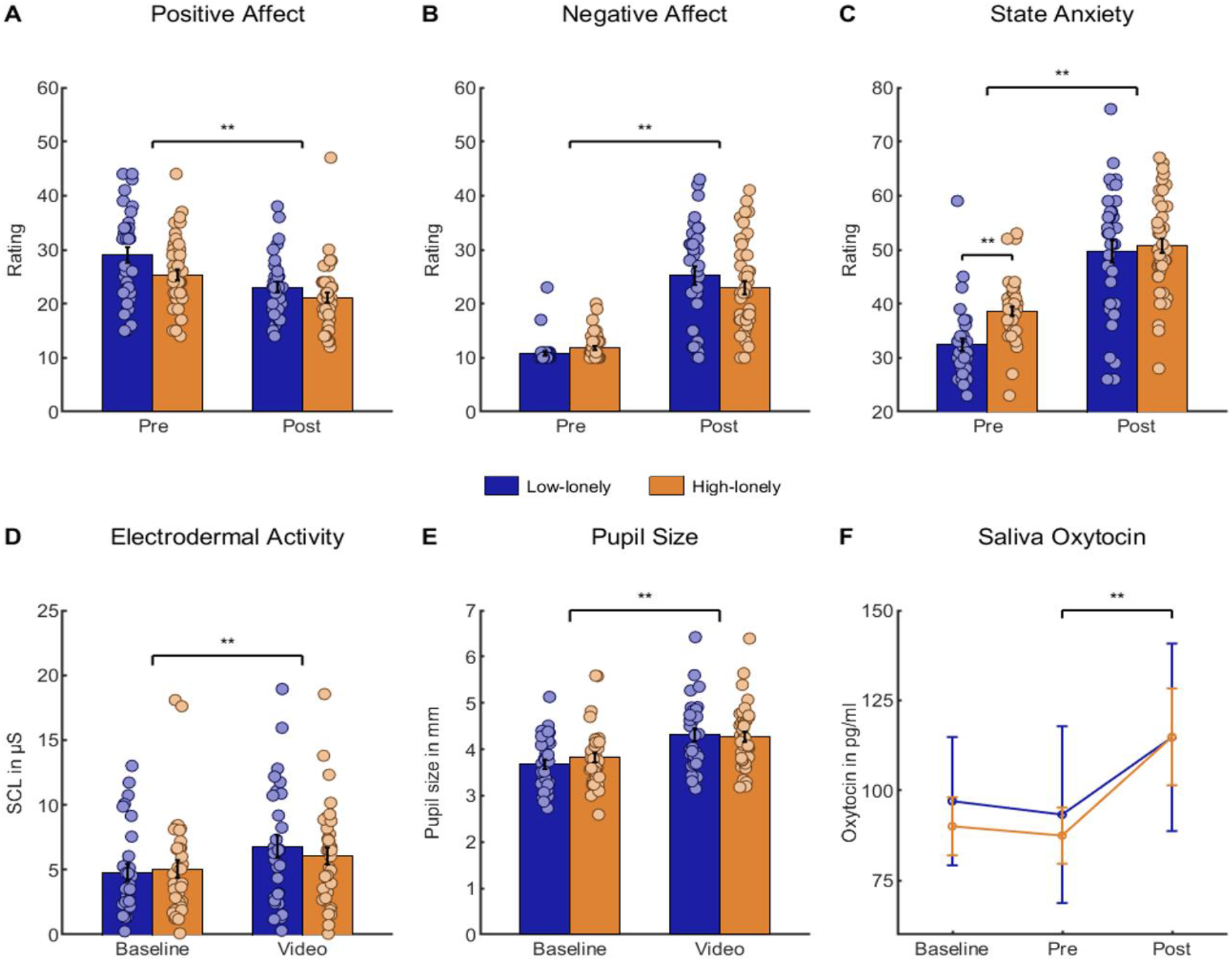
Acute psychosocial and physiological responses to the trauma paradigm were comparable across groups. Affect measured by the positive and negative affect schedule (PANAS) changed significantly, such that positive affect decreased (*t*_(75)_ = 8.13, *p* < 0.01, *d* = 0.74; **A**), while negative affect increased (*t*_(75)_ = 11.48, *p* < 0.01, *d* = 1.89; **B**). Baseline anxiety measured by the State Trait Anxiety Inventory (STAI) was increased in high-lonely subjects (*t*_(76)_ = 4.42, *p* < 0.01, *d* = 1.02; **C**) and increased across groups (*t*_(75)_ = 11.49, *p* < 0.01, *d* = 1.65). Physiological arousal was evident in increased skin conductance levels (*t*_(64)_ = 3.67, *p* < 0.01, *d* = 0.36; **D**) and pupil sizes (*F*_(1,65)_ = 133.96, *p* < 0.01, η_p_^2^ = 0.67; **E**) during the video. Furthermore, saliva oxytocin levels increased significantly after trauma exposure (*t*_(72)_ = 4.05, *p* < 0.01, *d* = 0.24; **F**). Error bars show the standard error of the mean (SEM). Abbreviations: pre, directly before the trauma paradigm; post, directly after the trauma paradigm; SCL, skin conductance level; ** *p* < 0.01.

Physiologically, there was an increase in the skin conductance level (main effect of time: *F*_(1,61)_ = 13.57, *p* < 0.01, η_p_^2^ = 0.18; shown in **Fig. 1D**) and pupil size (*F*_(1,65)_ = 133.96, *p* < 0.01, η_p_^2^ = 0.67; shown in **Fig. 1E**) compared to baseline. Furthermore, salivary oxytocin levels significantly increased after trauma exposure (*F*_(2,130)_ = 3.39, *p* = 0.04, η_p_^2^ = 0.05; post hoc *t*-test: *t*_(72)_ = 4.05, *p*_cor_ < 0.01, *d* = 0.47; shown in **Fig. 1F**). Thus, the trauma video elicited a psychological and physiological stress response regardless of sex and loneliness.

### Intrusive thoughts

Across loneliness groups, women experienced more intrusions than men (main effect of sex: *F*_(1,77)_ = 8.53, *p* = 0.01, η_p_^2^ = 0.10). However, our results revealed a significant interaction between loneliness and sex (*F*_(1,77)_ = 5.57, *p* = 0.02, η_p_^2^ = 0.07), such that loneliness was associated with more intrusive memories in men but fewer intrusions in women (shown in **Fig. 2A**). Post hoc *t*-tests further revealed that low-lonely women exhibited significantly more intrusions than low-lonely men (*t*_(33)_ = 3.97, *p_cor_* < 0.01, *d* = 1.39), while there was no significant sex difference in high-lonely individuals (*t*_(44)_ = 0.39, *p* = 0.70, *d* = 0.12). Furthermore, analysis of the desire to talk about the trauma movie yielded a pattern consistent with intrusion effects (*F*_(1,65)_ = 5.62, *p* = 0.02, η_p_^2^ = 0.08; shown in **Fig. 2B**). High-lonely woman showed a decreased desire, whereas high-lonely men exhibited an increased desire in contrast to low-lonely individuals. Again, post hoc *t*-tests revealed that low-lonely women showed an increased desire to talk in contrast to low-lonely men (*t*_(32)_ = 2.66, *p_cor_* = 0.046, *d* = 0.91). In addition, high-lonely subjects talked less about the movie (main effect of loneliness: *F*_(1,49)_ = 9.85, *p* < 0.01, η_p_^2^ = 0.17; shown in **Fig. 2C**), indicating that the sex-specific association of loneliness with the desire to talk about the traumatic experience did not lead to a similar pattern in actual trauma disclosure. Neither sex nor loneliness significantly affected intrusion stress ratings (all *p*s > 0.05).

**Fig. 2.**
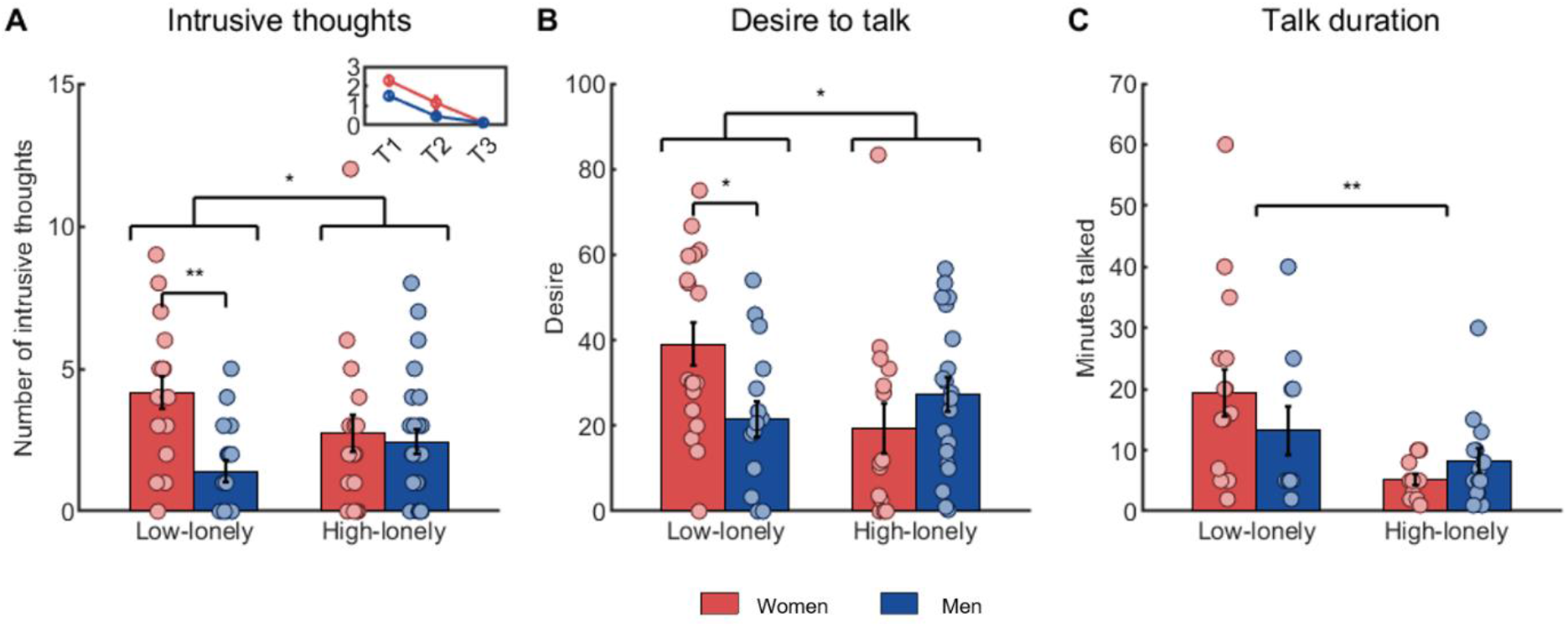
High-lonely men experienced more intrusions than low-lonely men in the three days following the trauma video, while this pattern was reversed in women (interaction effect: *F*_(1,77)_= 5.57, *p* = 0.02, η_p_^2^ = 0.07; **A**). The inlay shows the decrease in intrusions over the following three days. High-lonely men showed an increased desire to talk about the experience (from 0 = no desire to 100 = extreme desire) in contrast to low-lonely men. Women showed the reversed pattern (interaction effect: *F*_(1,65)_= 5.62, *p* = 0.02, η_p_^2^ = 0.08; **B**). Furthermore, high-lonely subjects talked less about their traumatic experience regardless of sex (main effect of loneliness: *F*_(1,49)_= 9.85, *p* < 0.01, η_p_^2^ = 0.17; **C**) Error bars show the standard error of the mean (SEM). Abbreviations: T1-T3, days after trauma exposure; * *p* < 0.05; ** *p* < 0.01.

### Emotional face-matching: fMRI effects

There was no significant interaction effect of sex and loneliness on the neural response to fearful faces per se, but amygdala habituation was characterized by sex*loneliness interactions. Habituation to fearful faces in the right amygdala was reduced in high-lonely men compared to high-lonely women, while this pattern was reversed in low-lonely individuals (interaction sex*loneliness: Montreal Neurological Institute (MNI)_xyz_: 34, 2, −22, *F*_(1,75)_ = 12.72, *p*_FWE_ = 0.04; Fearful _Block 1_ > Fearful _Block 3_; shown in **Fig. 3A**). Across groups, right amygdala habituation to fearful faces correlated negatively with the number of intrusions (*r*_(76)_ = −0.22 *p* = 0.049; Fearful _Block 1_ > Fearful _Block 3_).

**Fig. 3.**
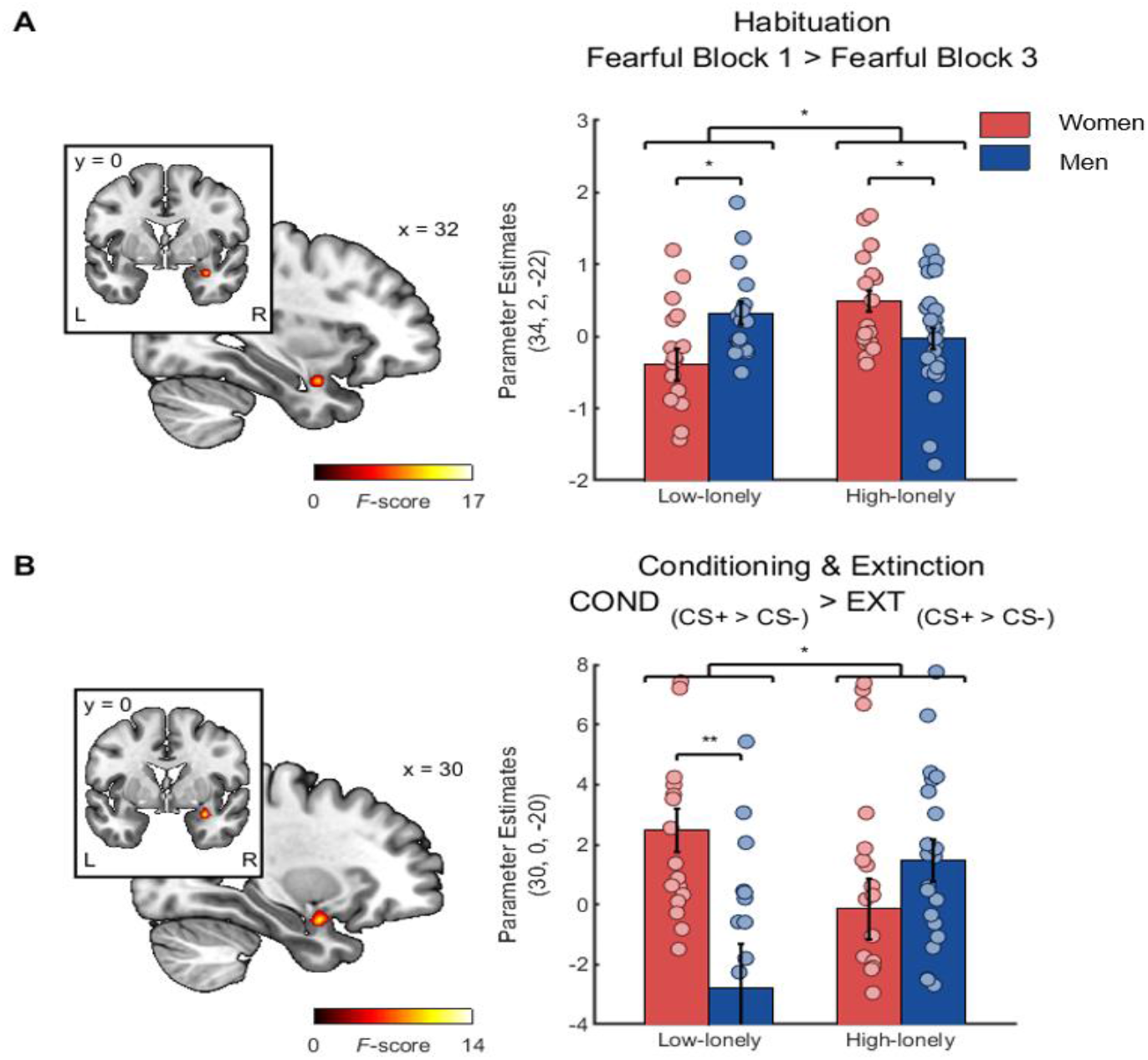
High-lonely men showed decreased right amygdala habituation (MNI_xyz_: 34, 2, −22, *F*_(1,75)_ = 12.72, *p*_FWE_ = 0.04; **A**) to fearful faces in contrast to high-lonely women and this pattern was reversed in low-lonely individuals. Likewise, high-lonely men exhibited stronger right amygdala activity (MNI_xyz_: 30, 0, −20, *F*_(1,72)_ = 12.62, *p*_FWE_ = 0.046; **B**) during fear conditioning than high-lonely women and this pattern was also reversed in low-lonely participants. Coordinates are in MNI space. Error bars show the standard error of the mean (SEM). Abbreviations: COND, conditioning; CS+, fear-associated conditioned stimulus; CS-, non-fear-associated conditioned stimulus; EXT, extinction; L, left; R, right; * *p* < 0.05; ** *p* < 0.01.

Further habituation analyses revealed a sex*loneliness interaction in functional connectivity. High-lonely men showed increased right amygdala coupling with the left superior parietal lobe in the habituation process to fearful faces (MNI_xyz_: −34, −52, −58, *k*_E(1,75)_ = 108, *p*_FWE_ = 0.01; Fearful _Block 1_ > Fearful _Block 3_; shown in **Fig. 4A**) in contrast to high-lonely women, while this pattern was reversed in low-lonely individuals. Further behavioral and neural results of the emotional face matching task are reported in the SI (shown in **Tab. S2/S3**).

**Fig. 4.**
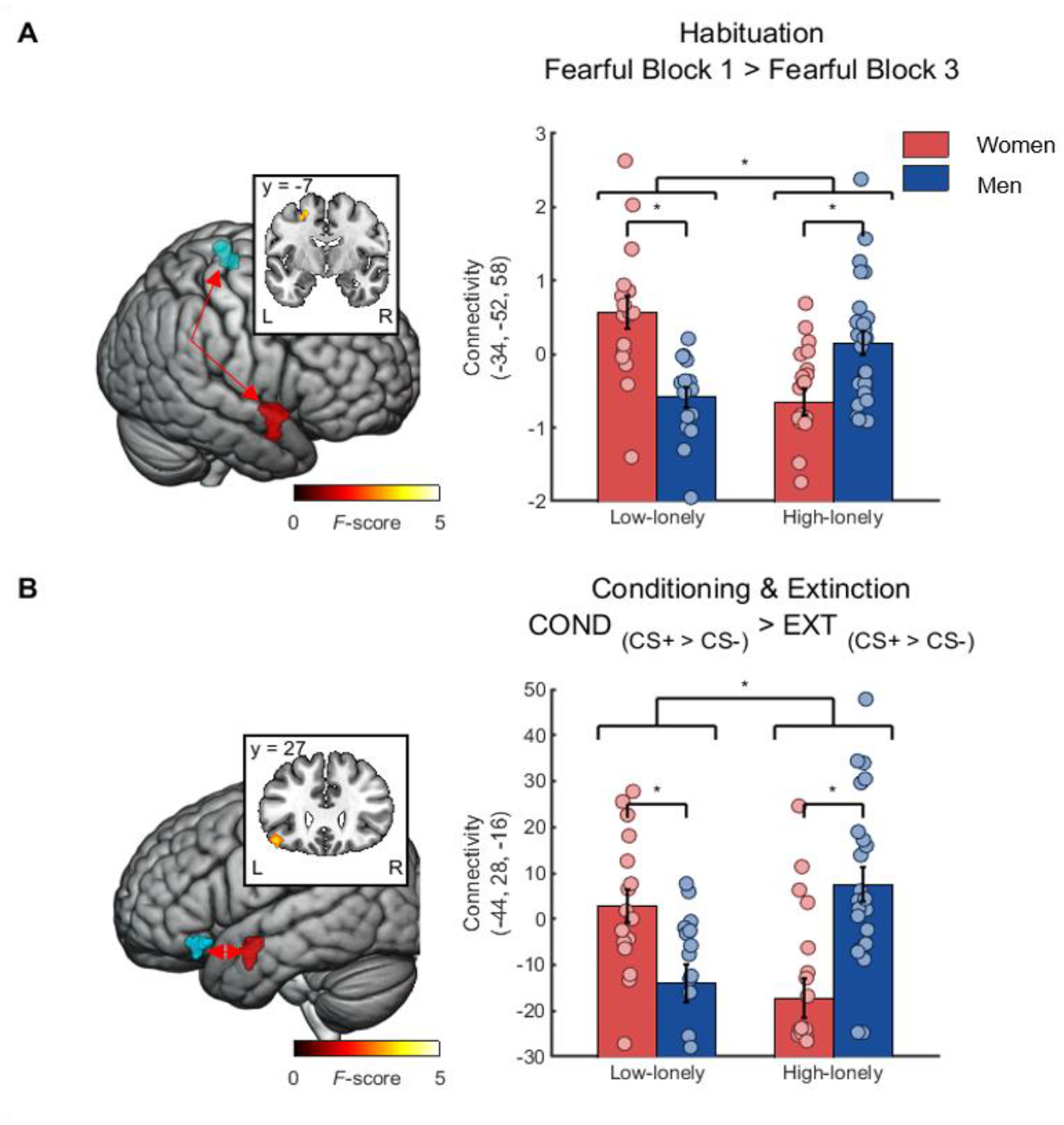
Increased coupling between the right amygdala (red cluster) as the seed region and left superior parietal lobule (blue cluster; MNI_xyz_: −34, −52, −58, *k*_E(1,75)_ = 108, *p*_FWE_ = 0.01; **A**) was observed during fear habituation in high-lonely men compared to high-lonely women, whereas this pattern was reversed in low-lonely individuals. Furthermore, connectivity between the left amygdala (red cluster) as the seed region and left orbitofrontal areas (blue cluster; MNI_xyz_: −44, 28, −16, *k*_E(1,75)_ = 98, *p*_FWE_ = 0.02; **B**) was increased during fear conditioning in high-lonely men compared to high-lonely women, and this pattern was reversed in low-lonely participants. Coordinates are in MNI space, and error bars show the standard error of the mean (SEM). Abbreviations: COND, conditioning; CS+, fear-associated conditioned stimulus; CS-, non-fear-associated conditioned stimulus; EXT, extinction; L, left; R, right; * *p* < 0.05.

### Fear conditioning and extinction: contingency ratings

Successful conditioning was evident in higher contingency ratings of the CS+ compared to the CS-in the second half of the COND task (interaction effect of time and condition: *F*_(1,64)_ = 54.79, *p* < 0.01, η_p_^2^ = 0.46). Likewise, a significant time*condition interaction (*F*_(1,63)_ = 49.23, *p* < 0.01, η_p_^2^ = 0.44) for the contingency ratings showed reduced shock expectations in the course of the EXT task (shown in **Tab. S4**). In addition, a time*condition interaction with sex and loneliness was evident such that high-lonely men showed higher contingency ratings (i.e., expected more electric shocks) to the CS+ in the second half of the COND phase than high-lonely women (time*condition*sex*loneliness; *F*_(1,64)_ = 5.41, *p* = 0.02, η_p_^2^ = 0.08). There were no significant interactions with the factor sociality or sex * loneliness interactions in the EXT phase.

### Fear conditioning and extinction: fMRI effects

In the COND phase, the CS+ elicited activations in a fear conditioning network [37] including the amygdala (COND _CS+ > CS-_ MNI coordinates and statistics are listed in **Tab. S5**). Importantly, amygdala reactivity to fear signals in the early phase of COND compared to that in EXT was associated with loneliness in a sex-specific manner (sex*loneliness interactions: MNI_xyz_: 30, 0, −20, *F*_(1,72)_ = 12.62, *p*_FWE_ = 0.046; COND _CS+ > CS-_ > EXT _CS+ > CS-_; shown in **Fig. 3B**). This effect was driven by a sex*loneliness interaction in the COND phase (MNI_xyz_: 30, 4, −20, *F*_(1,72)_ = 14.37, *p*_FWE_ = 0.02; COND _CS+ > CS-_). High-lonely men exhibited higher amygdala activation than high-lonely women, while this effect was reversed in low-lonely individuals.

We also observed a loneliness*sex interaction in the functional connectivity of the amygdala during fear conditioning/extinction. High-lonely men exhibited a stronger coupling between the left amygdala and orbitofrontal cortex (MNI_xyz_: −44, 28, −16, *k*_E(1,72)_ = 98, *p*_FWE_ = 0.02; COND _CS+ > CS-_ > EXT _CS+ > CS-_; shown in **Fig. 4B**) compared to high-lonely women during the conditioning of fear signals and this pattern was reversed for low-lonely individuals.

Importantly, including psychiatric symptoms that differed between groups (shown in **Tab. 1 and Tab. S6**), social support, hormonal contraception, and estradiol levels as covariates did not change the significant sex*loneliness interactions observed for intrusions and parameter estimates of significant clusters (see SI). Taken together, these findings indicate that fear habituation and conditioning mechanisms in an amygdala network vary as a function of loneliness and sex.

## Discussion

The present study aimed to probe loneliness as a risk factor for increased physiological and psychological responses to an experimental trauma film. We further examined whether loneliness effects were moderated by sex and related to changes in the neural processing of fear signals. Our results revealed a significant interaction between sex and loneliness in intrusive thought formation such that loneliness was positively associated with more intrusions in men, but not women. A similar pattern emerged at the neural level, with both reduced amygdala habituation to repeated fearful faces and amygdala hyperreactivity during the conditioning of fear signals in high lonely men, but not in women. Our findings indicate that loneliness is indeed a risk factor for increased intrusions after trauma exposure, and this relates to amygdala reactivity in a network responsible for fear conditioning and habituation in high-lonely men.

The experimental trauma paradigm elicited marked stress responses evident in significant psychological and physiological changes across all subjects. We did not observe significant loneliness or sex effects on these acute responses after the trauma film, indicating that loneliness may be more important in long-term coping with the traumatic experience. As expected, high-lonely subjects talked less about their traumatic experience than low-lonely individuals. Disclosure of emotional and traumatic events is known to reduce distress and may promote extinction of fear-related memories [48, 49]. Furthermore, discussing traumatic memories reduces PTSD symptoms, and delayed disclosure predicts PTSD development [50–53]. Interestingly, reduced trauma disclosure cannot completely explain the loneliness-associated increase in intrusive thoughts observed in men, because high-lonely women also reported less trauma disclosure and experienced fewer intrusions than low-lonely women. In the current sample, in contrast to men, high-lonely women may be less vulnerable to trauma-induced intrusions since they also indicated less desire to talk about the trauma film relative to low-lonely women. Therefore, in the same way that loneliness results from a discrepancy between desired and actual social connectedness, a mismatch between the desired and achieved trauma disclosure may be particularly important for individuals to cope with intrusive thoughts.

The amygdala is a well-known processing hub of fear-related stimuli [54–56] and amygdala hyperreactivity is a risk factor for as well as a consequence of trauma-related disorders [57–60]. Sex-differences in amygdala lateralization and habituation have been previously observed, with women exhibiting more activity in the left hemisphere related to the subsequent memory for emotionally arousing images and showing more persistent bilateral amygdala responses to negative stimuli than men [30, 61, 62]. Intriguingly, low-lonely women showed significantly less amygdala habituation and experienced significantly more intrusions than low-lonely men and increased amygdala habituation correlated with reduced intrusions across groups. Our findings are consistent with previous studies that showed that decreased amygdala habituation is associated with heightened anxiety levels [63–67] and PTSD symptom severity [68]. Furthermore, increased functional connectivity between the amygdala and the superior parietal lobe in high-lonely men may constitute a prospective risk factor for heightened intrusive thoughts since the parietal lobe is part of a common network responsible for intrusive thought formation [69]. Moreover, PTSD patients exhibit increased parietal activations during script-driven trauma imagery leading to dissociative responses [70].

Furthermore, high-lonely men exhibited heightened amygdala responses to the CS+ and functional connectivity with the orbitofrontal cortex during conditioning compared to low-lonely men. Both amygdala and orbitofrontal cortex activity have been frequently linked to CS+/CS-differentiation during fear learning [71]. The loneliness-related amygdala activation changes during habituation and conditioning in men were evident across social and non-social stimuli, which is in line with previous studies suggesting that loneliness fosters hypervigilance for threat cues [7, 72, 73]. The absence of this effect in high-lonely women could be driven by hormonal factors with high-lonely women showing increased estradiol levels compared to low-lonely women in our sample. Estradiol administration improved extinction recall after fear extinction [74, 75], and low levels of estradiol in women were linked to increased fear network responses to trauma films [76]. However, the observed sex differences cannot be completely explained by hormonal factors either because women reported more intrusions across loneliness groups despite having higher estradiol levels than men. It is conceivable that the content of the trauma film was more distressing for women than men, but we did not detect significant sex differences in the acute stress responses, and a previous study found no evidence for an interaction between sex and intrusive memories induced by different trauma films [77]. The unwillingness of men to admit loneliness and higher stigmatization of men who express feelings of loneliness [78, 79] might have contributed to the observed sex differences. Taken together, our data suggest that loneliness has a sex-specific impact on the way threat cues are processed during fear conditioning and fear habituation.

The present study had several limitations. First, our sample consisted of women with and without hormonal contraception. Although we did control for the use of hormonal contraception and measured hormonal blood levels to control for menstrual cycle-related hormone changes, future studies are warranted to further delineate the hormonal basis of sex differences in the effects of loneliness. Second, the experimental trauma paradigm is widely used and well established to explore the neurobiological mechanisms underlying acute and prolonged trauma responses, but further clinical studies in a real-life setting are required to gauge whether our findings can be extrapolated to patients with trauma exposure.

Collectively, our results provide evidence that loneliness may confer vulnerability to increased intrusive thoughts in men following an experimental trauma. In addition, high-lonely men were characterized by an increased desire to talk about the trauma film and reduced actual trauma disclosure. This phenotype relates to altered limbic processing driven by amygdala hyperreactivity during fear conditioning and habituation. Based on these findings, secondary prevention strategies should take sex differences in loneliness into account and focus on improving the social connectedness of high-lonely men to mitigate the sequelae of traumatic experiences.

## Statements

### Study approval statement

This study protocol was reviewed and approved by the institutional review board of the medical faculty of the University of Bonn [Approval number: 248/16] and carried out in compliance with the latest revision of the Declaration of Helsinki.

### Consent to participate statement

Written informed consent was obtained from all participants included in this study.

### Author contributions

M.M., and D.S. designed the experiment; M.M., J.N., D.S. and J.D. conducted the experiments; M.M., B.S-W. and D.S. analyzed the data. M.M. and D.S. wrote the manuscript. All authors read and approved the manuscript in its current version.

### Data availability statement

Data analysis was preregistered prior to conducting any analyses and behavioral and parameter estimate data will be publicly available after publication in the open science framework (https://osf.io/np9wr/?view_only=d1cf3677f9ea42578eaceb882069ce89). Second level fMRI data will be publicly available at NeuroVault (https://neurovault.org/collections/HOUZNUPY/).

