## Supplementary_Information for "Lonely in the dark: trauma memory and sex-specific dysregulation of amygdala reactivity to fear signals"

**Supplementary Methods**

**Power analysis**

To the best of our knowledge, no study has examined the association between loneliness and individuals’ responses to viewing a trauma film or fear conditioning/extinction and fear habituation. Thus, we used G*Power 3 to conduct an a-priori power analysis for the project based on the effect size obtained in a functional magnetic resonance imaging (fMRI) study investigating the neural processing of social stimuli as a function of perceived social isolation [1]. Cacioppo et al. observed a correlation of *r* = -.46 for the reactivity of the ventral striatum to positive social stimuli with the UCLA loneliness scores of participants. To reliably replicate this effect of loneliness on the neural processing of social stimuli (with α = 0.05 and power = 0.99), at least 71 participants had to be evaluated. To account for possible dropouts, we planned to assess at least 80 participants.

**Online recruiting**

We used the UCLA loneliness scale (LS) as an online questionnaire to recruit eligible subjects for the study. Out of 4515 participants, 97 subjects were invited to a screening session to evaluate the inclusion criteria: LS score above or equal to 50 (high-lonely) or 25 or below (low-lonely), aged 18-65 years, no current physical or psychiatric disorder as assessed via self-disclosure and the Mini-International Neuropsychiatric Interview [2], no psychotherapy, no current psychotropic medication, no illicit drug use in the previous four weeks, and eligibility for magnetic resonance imaging scanning. Subjects were screened prior to the testing session. Fifteen participants had to be excluded after the screening session because they failed to fulfil the inclusion criteria.

**Questionnaires**

Prior to the screening session, subjects completed an online questionnaire consisting of personal data and the UCLA LS. Subjects with scores above 50 and below 25 were invited for screening sessions. Screenings consisted of interviews about their medical history and the Mini-International Neuropsychiatric Interview [2]. Furthermore, we assessed alexithymia (Toronto Alexithymia Scale [TAS]) [3], perceived stress (Perceived Stress Scale [PSS-10]) [4], perceived social support (Fragebogen zur Sozialen Unterstützung, short version K-14 [F-SozU]) [5], social interaction anxiety (Liebowitz Social Anxiety Scale [LSAS]) [6], social network size (Social Network Size Questionnaire [SNS]) [7], childhood trauma (childhood trauma questionnaire [CTQ]) [8], depression symptoms (Beck Depression Inventory [BDI]) [9], and trait anxiety (State Trait Anxiety Inventory [STAI]) [10]. Before and directly after participants viewed the trauma video, we assessed positive and negative affect (positive and negative affect schedule [PANAS]) [11], as well as arousal and valence ratings. In addition, dissociative symptoms (Dissociative Symptoms Scale [DSS]) [12] were measured after participants viewed the trauma video. All questionnaires were presented with Qualtrics software (Provo, USA).

**Emotional face-matching task**

The trial duration was 5 s with a 10 s pause interblock interval in which a fixation-cross was displayed. Participants were asked to react to the presented stimuli as quickly as possible.

**Fear conditioning and extinction tasks**

To identify a stimulation intensity that was uncomfortable, but not painful, participants rated different intensities beforehand on a scale from 0 to 100 (0 = not uncomfortable; 100 = most uncomfortable feeling imaginable). The stimulation intensity was set to reflect a rating of 60. Stimulation intensity was increased stepwise until participants first reached a rating of 60. To further validate this result, intensity was then lowered twice by two intensity steps, followed by the original intensity. If ratings were comparable, the subject received three shocks for habituation. If the ratings differed from the original rating, the intensity was again increased stepwise followed by the adaptive process until a rating of 60 was reached. The trials were interleaved with an interstimulus interval (ISI) that was jittered between 5 s and 7 s (mean: 6 s). After the COND phase, participants were informed that there would be another round of the same experiment.

**Neuroendocrine parameters and analysis**

Saliva samples collected for oxytocin measurement were acquired using commercial sampling devices (Salivettes, Sarstedt, Germany) and were cooled directly after collection. Samples were centrifuged at 4,000 rpm for 2 min and stored at -80°C until assayed. Before the fMRI session, blood samples were collected to measure estradiol, testosterone, progesterone, luteinizing hormone (LH), follicle-stimulating hormone (FSH), and dehydroepiandrosterone (DHEAS). Estradiol and testosterone were analyzed in line with the manufacturer’s instructions (Siemens Healthineers, Eschborn, Germany) and by fully automated homogeneous sandwich chemiluminescent immunoassays based on LOCI™ technology on a Dimension Vista™ system. For testosterone, the detection limit of the assay was 0.025 ng/ml and the coefficients of variation for intra-assay and inter-assay precision were 4.7% and 6.7%, respectively. Estradiol was tested with a detection limit of 5 pg/ml and the intra-assay and inter-assay precision variation coefficients were 5.5% and 5.9%, respectively. Serum progesterone was analyzed by applying a fully automated solid-phase competitive chemiluminescent enzyme immunoassay on an Immulite™ 2000xpi system according to the manufacturer´s instructions (Siemens Healthineers) with a detection limit of 0.1 ng/ml. Intra-assay and the inter-assay precision varied between 4.2% and 5.5%. Serum LH, FSH, and DHEAS were analyzed by fully automated electrochemiluminescence immunoassays (ECLIA, Elecsys tests) on a Cobas e801 analyzer (Roche Diagnostics, Mannheim, Germany) according to the manufacturer´s instructions (Roche Diagnostics). The coefficients of variation for intra-assay and inter-assay precision were 1.63% and 2.06% for LH, 2.37% and 2.71% for FSH, and 2.28% and 2.47% for DHEAS, respectively. There was minimal cross-reactivity of all assays with other related compounds. Saliva oxytocin was analyzed with an ELISA kit (ENZO Life Sciences GmbH, Lörrach, Germany) with a detection limit of 15 pg/ml. Intra- and inter-assay precision varied between 7.4% and 11.22%.

**FMRI data acquisition**

All fMRI data were acquired using a 3T Siemens TRIO MRI system (Siemens AG, Erlangen, Germany) with a Siemens 32-channel head coil. Functional data were obtained using a T2*-weighted echoplanar (EPI) sequence [repetition time (TR) = 2690 ms, echo time (TE) = 30 ms, ascending slicing, matrix size: 96 x 96, voxel size: 2 x 2 x 3 mm³, slice thickness = 3.0 mm, distance factor = 10%, field of view (FoV) = 192 mm, flip angle 90°, and 41 axial slices]. High-resolution T1-weighted structural images were collected on the same scanner (TR = 1660 ms, TE = 2.54 ms, matrix size: 256 x 256, voxel size: 0.8 x 0.8 x 0.8 mm³, slice thickness = 0.8 mm, FoV = 256 mm, flip angle = 9°, 208 sagittal slices). To control for inhomogeneity of the magnetic field, field maps were obtained for each T2*-weighted EPI sequence and were included during preprocessing of the fMRI data (TR = 392 ms, TE [1] = 4.92, TE [2] = 7.38, matrix size: 64 x 64, voxel size: 3 x 3 x 3, slice thickness = 3.0 mm, distance factor = 10%, FoV = 192 mm, flip angle 60°, 37 axial slices). In both fMRI tasks, stimuli were presented on a 32-inch MRI compatible TFT LCD monitor (NordicNeuroLab, Bergen, Norway) placed at the rear end of the magnet bore. Participants could choose their responses with an MRI-compatible response grip system (NordicNeuroLab AS, Bergen, Norway). The paradigms were written in Presentation code (Neurobehavioral Systems, Inc., Berkeley, USA, www.neurobs.com). High-resolution anatomical images were acquired after the functional images.

**FMRI data preprocessing**

FMRI data were preprocessed and analyzed using standard procedures in SPM12 (Wellcome Trust Center for Neuroimaging, London, UK; http://www.fil.ion.ucl.ac.uk/spm) implemented in MATLAB (The MathWorks Inc., Natick, MA). The first five volumes of each functional time series were discarded to allow for T1 equilibration. Functional images were corrected for head movements between scans by an affine registration. Images were initially realigned to the first image of the time series before being re-realigned to the mean of all images. To correct for signal distortion based on B0-field inhomogeneity, the images were unwarped by applying the voxel displacement map (VDM file) to the EPI time series (Realign & Unwarp). Normalization parameters were determined by segmentation and non-linear warping of the structural scan to reference tissue probability maps in Montreal Neurological Institute (MNI) space. Normalization parameters were then applied to all functional images which were resampled at 2 x 2 x 2 mm³ voxel size. For spatial smoothing, a 6-mm full-width at half-maximum (FWHM) Gaussian kernel was used. Raw time series were detrended using a high-pass filter (cut-off period, 128 s).

**Emotional face-matching: fMRI analyses**

For first level analyses, the four conditions (happy, fearful, and neutral faces, and houses) were modeled by a boxcar function convolved with a hemodynamic response function. Furthermore, we assessed habituation by calculating the mean response amplitude differences between the first and third blocks for each condition on the first level. Button presses were included as regressors of no interest. Movement parameters were entered as confounding regressors in the design matrix using the artifact detection toolbox (ART, https://www.nitrc.org/projects/artifact_detect, RRID: SCR_005994). Any subjects with >20% volumes identified as outliers (> 1.5 mm/°) by ART were excluded. In total, one participant had to be excluded due to technical errors, and two participants had to be excluded due to excessive head motion resulting in a final sample of 79 subjects. On the second level, the main contrasts of interest were compared between groups of participants using a full factorial model with the two factors loneliness (high-lonely vs. low-lonely) and sex (women vs men). Button presses were included as regressors of no interest. Based on our hypotheses, the analysis was conducted using the anatomically defined regions-of-interest (ROIs) of the amygdala, anterior cingulate cortex (ACC), insular cortex, and nucleus accumbens derived from the WFU PickAtlas (https://www.nitrc.org/projects/wfu_pickatlas/, RRID: SCR_007378). The significance threshold for these ROI analyses was set to *p* < 0.05, familywise error-corrected (p_FWE_) for multiple comparisons based on the size of the ROI. Parameter estimates of significant contrasts were extracted using MarsBar (https://www.nitrc.org/projects/marsbar, RRID: SCR_009605) and further analyzed in SPSS 25 (IBM Corp., Armonk, NY, USA). Furthermore, an exploratory whole-brain analysis was performed to detect task effects (cluster defining threshold *p* < 0.001; significance threshold *p*_FWE_ < 0.05 corrected at peak level). In addition, generalized psychophysiological interaction (gPPI) analysis was conducted using the CONN toolbox 18.a (www.nitrc.org/projects/conn, RRID:SCR_009550) [13] with the same preprocessed data, ROIs, regressors, and contrasts that were used in the SPM analyses. After denoising, the first levels for each subject were calculated using the psychological (task effect) and physiological factors (BOLD time series). Bivariate regression was used to measure the task specific connectivity compared to the implicit baseline. Mixed-design ANOVAs were used to examine task-specific connectivity main and interaction effects of loneliness and sex. A height threshold of *p* < 0.001 was used as a cluster-forming threshold to define significant clusters. Beta weights of significant effects of interest were extracted using MarsBar and further analyzed in SPSS.

**Fear conditioning and extinction: behavioral analyses**

The reaction times (RTs) of contingency ratings were assessed for all trials in which the rating occurred 4 s after stimulus onset (before the electric impulse).

**Fear conditioning and extinction: fMRI analyses**

For the conditioning (COND) /extinction (EXT) paradigm, a two-stage approach based on the general linear model implemented in SPM12 was used for statistical analyses. On the first level, participants’ individual data were modeled using a fixed-effect model. Onsets and durations of the six experimental conditions (‘COND’, ‘EXT’, ‘social’, ‘non-social’, ‘CS+’, and ‘CS-‘) were modeled by a stick function convolved with a hemodynamic response function (HRF). Movement parameters were included in the design matrix as confounds using ART. Any subjects with >20% volumes identified as outliers (> 1.5 mm/°) by ART were excluded, resulting in a final sample size of 76 individuals (three subjects were excluded from the analysis due to technical errors and three subjects due to excessive head motion). Respiratory data were used as confound regressors created using the MATLAB PhysIO Toolbox (<https://www.nitrc.org/projects/physio>) [14]. On the second level, the main contrasts of interest were computed using a full factorial model with the two factors loneliness (high-lonely vs. low-lonely) and sex (women vs. men). Button presses and electrical shocks were included as regressors of no-interest. Analysis was conducted using the anatomically defined amygdala, insular cortex, ACC, posterior cingulate cortex (PCC), medial prefrontal cortex (mOFC), and hippocampus as ROIs, according to the WFU PickAtlas. All ROIs were derived from recent meta-analyses of fMRI COND/EXT experiments [15, 16]. The significance threshold for these ROI analyses was set to *p* < 0.05, familywise error-corrected (*p*_FWE_) for multiple comparisons based on the size of the ROI. Parameter estimates of significant contrasts were extracted using MarsBar and further analyzed in SPSS 25. In addition, an exploratory whole-brain analysis was performed to detect task effects (cluster defining threshold *p* < 0.001; significance threshold *p*_FWE_ < 0.05 corrected at peak level). Furthermore, exploratory whole-brain and ROI analyses were performed using intrusive thoughts as a covariate. To further examine the potential influence of sex and loneliness on task-based functional connectivity, a gPPI analysis was conducted. The analysis was operated with the same preprocessed data, ROIs, regressors and contrasts that were used in the SPM analyses. Task-based functional connectivity was analyzed using the CONN toolbox. First and second levels were calculated as mentioned in the emotional face-matching analyses using the same height and cluster forming thresholds. Beta weights of significant clusters were extracted using MarsBar and further analyzed in SPSS.

**Fear conditioning and extinction: physiological data assessment**

Physiological responses during the COND/EXT tasks were measured with a Biopac MP150 system. Electrodermal activity (EDA) was measured at a sampling rate of 1000 Hz from Ag/AgCl electrodes filled with isotonic electrolyte gel on the thenar and hypothenar of the left hand. Respiration was measured by a TSD221-MRI transducer (MP150, Biopac Systems Inc., Goleta, USA). EDA data were preprocessed and analyzed with Acqknowledge 4.3 software (Biopac Systems Inc., Goleta, USA). The EDA data were smoothed (median value smoothing factor: 63) and a low-pass filter (frequency cutoff 1 Hz) was applied. The remaining non-physiological artifacts were removed by visual inspection. Phasic components were derived from the tonic EDA before the skin conductance responses (SCR) were measured. SCRs were measured in a time window between 0.5 and 4.5 s after stimulus presentation. A SCR was defined as a change of at least 0.01 µS. Prior to data analysis, a square root transformation was applied to the SCR amplitudes. The magnitudes of SCRs were further analyzed and compared between groups using SPSS 25.

**Experimental trauma paradigm: physiological data assessment**

Pupil sizes were measured with an eye-tracking system. Participants were seated in front of a Tobii TX300 binocular eye-tracker (Tobii AB, Danderyd, Sweden) with a 23-inch display. The Tobii TX300 binocular eye-tracker had a maximum resolution of 1920 x 1080 pixels, 0.01° precision, and a sampling rate of 300 Hz. Participants’ eye movements were calibrated prior to the experimental trials. Pupil sizes were measured with the Tobii Studio eye-tracking software version 3.2.3. After the calibration procedure, participants were presented with a 40-s neutral scene of the movie to obtain a baseline measure of the physiological data (pupil size and EDA). EDA data were measured with a Biopac MP150 system electrodes filled with isotonic electrolyte gel on the thenar and hypothenar of the left hand at a sampling rate of 1000 Hz from Ag/AgCl. EDA data were preprocessed as described above. Phasic components were derived from tonic EDA before the skin conductance levels (SCLs) were assessed. SCLs were measured in µS. For the trauma film, we used a 24-minute-long movie clip derived from the movie “I Spit on Your Grave”.

**Online diaries**

The participants completed online intrusion diaries at home in the evening for three consecutive days after trauma exposure. In the intrusion diary, the participants stated the number of intrusions (defined as involuntary recollections relating to film events that appear, apparently spontaneously, in consciousness) and rated the distress caused by these intrusions on a visual analogue scale ranging from 0 (no distress) to 100 (extreme distress). Furthermore, participants were asked whether and how long they talked about the trauma video and about their desire (0 = no desire to 100 = strong desire) for trauma disclosure.

**Further statistical analyses**

Hormonal blood parameters were analyzed using standard procedures including ANOVAs with the between-subject factors sex and loneliness and Bonferroni corrected post-hoc *t*-tests. Mixed-design ANOVAs were calculated for the emotional face-matching task for each condition with the between-subject factors sex and loneliness to examine differences in RTs and correct response rates (CRs). Habituation effects in RTs of the emotional face matching task were tested with mixed-design ANOVAs with the additional within-subject factor time “Block _1_ vs. Block _3_”. RTs and CRs were further analyzed with Bonferroni corrected post hoc *t*-tests if necessary. Likewise, mixed-design ANOVAs were used to test for changes in RTs and SCRs in the fear conditioning and extinction paradigm with the within-subject factors task “COND vs. EXT”, “CS+ vs. CS-“, and the between-subject factors sex and loneliness. Additional mixed-design ANOVAs included the between-subject factors of sociality denoted as “social vs. non-social” and time, defined as “first half vs. second half”. If the assumption of sphericity was significantly violated as assessed by Mauchly’s tests, Greenhouse Geisser corrections were applied. Partial eta-squared and Cohen’s *d* were calculated as measures of effect size.

**Missing values**

Due to technical errors (n = 3) or excessive head motion (n = 3) six subjects had to be excluded from the conditioning and extinction paradigm. In addition, three subjects had to be excluded from the emotional face-matching task due to technical errors (n = 1) or excessive head motion (n = 2). In total, 18 out of 246 online diaries were missing resulting in a data loss of 7.32%. Due to connection issues, four pre and four post video questionnaires about affect and state anxiety were lost. In the analysis of physiological reactions to the trauma video, 11 eye tracking datasets and 16 EDA datasets were lost due to technical errors or artifacts. Furthermore, eight blood samples were lost because of problems with sample assessment or analysis. Samples with missing values or concentrations below detection limits were discarded from the analysis (estradiol: n = 13; testosterone: n = 13; progesterone: n = 22; LH: n = 16; FSH: n = 13; SHBG: n = 33; DHEAS: n = 9).

**Supplementary Results**

**Additional analysis for the pupil responses to the trauma video**

We additionally analyzed pupil sizes separately for each eye and found a significant increase in size for the left (main effect of time: *F*_(1,66)_ = 98.13, *p* < 0.01, η_p_^2^ = 0.60) and right pupils (main effect of time: *F*_(1,65)_ = 147.94, *p* < 0.01, η_p_^2^ = 0.70). However, there were no significant main or interaction effects of sex and loneliness.

**Control analyses**

Inclusion of psychiatric symptoms (i.e., depressive symptoms, alexithymia, social and trait anxiety, childhood maltreatment, and perceived stress), social support, social network quality, and use of hormonal contraception as separate covariates did not change the significant interaction between sex and loneliness observed for intrusions and the desire to talk (all interactions *p*s < 0.05). Likewise, inclusion of estradiol blood concentrations as a covariate did not change the significant interaction between sex and loneliness observed for intrusions, however, the interaction effect on the desire to talk was no longer significant when estradiol concentration was included as covariate (*p* > 0.05). Furthermore, including the same variables as covariates in the mixed-design ANOVAs of parameter estimates of significant clusters did not change the observed significant sex*loneliness interactions.

**Hormonal blood parameters**

Blood samples were collected to measure testosterone, progesterone, estradiol, DHEAS, SHBG, LH and FSH concentrations. High-lonely women exhibited higher estradiol levels than low-lonely women at the fMRI session (*t*_(16.55)_ = 2.62, *p**_cor_* = 0.04, *d* = 0.87). This effect was not significant in the subsample of women not using hormonal contraception (*t*_(13.39)_ = 2.74, *p_cor_* = 0.08, *d* = 0.97). Blood concentrations for each group are shown in the supplementary information (cf. **Tab. S1**).

**Emotional face matching: reaction times**

Men showed significantly slower RTs to fearful faces compared to women across task blocks (main effect sex: *F*_(1,78)_ = 4.29, *p* = 0.04, η_p_^2^ = 0.05). No differences in CRs were observed between groups of participants (all *p*s > 0.05). Analyses of RT habituation revealed a significant effect of time (*F*_(1.69,124.67)_ = 4.54, *p* = 0.02, η_p_^2^ = 0.06) showing that subjects reacted faster in the last block. Across blocks, a significant loneliness * sociality was evident (*F*_(1.95,144.38)_ = 3.17, *p* = 0.05, η_p_^2^ = 0.04), but post hoc tests revealed no significant differences between high-lonely and low-lonely individuals (all *p*s > 0.05). A main effect of sociality (*F*_(1.95,144.38)_ = 7.81, *p* < 0.01, η_p_^2^ = 0.10) indicated that subjects reacted faster to face stimuli than to house stimuli (happy: *t*_(78)_ = 2.71, *p_cor_* = 0.048, *d* = 0.30, fearful: *t*_(78)_ = 6.35, *p_cor_* < 0.01, *d* = 0.71, neutral: *t*_(78)_ = 3.16, *p_cor_* = 0.01, *d* = 0.37). Likewise, subjects reacted faster to fearful faces than to happy (*t*_(78)_ = 4.75, *p_cor_* < 0.01, *d* = 0.53) and neutral faces (*t*_(78)_ = 3.59, *p_cor_* = 0.01, *d* = 0.40). There were no sex and loneliness interactions (all *p*s > 0.05). For RTs and CRs see **Tab. S2**.

**Emotional face-matching: fMRI effects**

Across groups, whole-brain analysis showed increased activity to face stimuli compared to non-social stimuli (i.e., houses) in a network including the hippocampus, amygdala, and frontal regions (Faces > Houses). Furthermore, middle temporal gyrus activity was increased in the contrast fearful faces larger neutral faces (Fearful > Neutral; MNI coordinates and cluster sizes are listed in **Tab.** **S7**). Additional ROI analysis showed a main effect of loneliness in the comparison between all face stimuli and non-social stimuli, with high-lonely subjects showing a decreased activity in the right ACC (MNI_xyz_: 16, 28, 24, *F*_(1,75)_ = 16.40, *p*_FWE_ = 0.04; Faces > Houses). Furthermore, high-lonely subjects showed increased activity in the left insula in response to fearful faces compared to low-lonely participants (MNI_xyz_: -34, 14, 0, *F*_(1,75)_ = 17.52, *p*_FWE_ = 0.04; Fearful > Neutral). No significant sex effects or sex*loneliness interactions were observed in these contrasts (all *p*s > 0.05).

Whole-brain analysis of habituation effects revealed decreased activity in response to the repeated presentation of face stimuli compared to non-social stimuli (i.e. houses) in the cuneus, lingual, and fusiform gyrus (Faces _Block 1 > Block 3_ > Houses _Block 1 > Block 3_). Habituation effects on fearful faces were evident in the superior frontal gyrus and supramarginal gyrus (Fearful _Block_ _1_ > Fearful _Block 3;_ MNI coordinates and cluster sizes are shown in **Tab.** **S3**). Furthermore, ROI analysis of task effects revealed right amygdala (MNI_xyz_: 18, -2, -14, *t*_(78)_ = 3.21, *p*_FWE_ = 0.05; Fearful _Block 1_ > Fearful _Block 3_) habituation to fearful faces and habituation to all faces in the left amygdala (MNI_xyz_: -26, 4, -18, *t*_(78)_ = 3.25, *p*_FWE_ = 0.04; Faces _Block 1_ > Faces _Block 3_).

We also observed a significant sex*loneliness interaction for the left amygdala habituation to all faces which was reduced in high-lonely women compared to high-lonely men and the opposite pattern was evident in low-lonely individuals (MNI_xyz_: -30, -2, -22, *F*_(1,75)_ = 17.53, *p*_FWE_ = 0.01; Faces _Block 1_ > Faces _Block 3_). Collectively, amygdala habituation and functional connectivity in high-lonely men seemed to be most pronounced in response to fearful stimuli, whereas amygdala habituation in high-lonely women seemed to be altered regardless of the emotional valence of the social stimuli.

Additionally, we found a significant sex*loneliness interaction for the habituation to all face stimuli in the right insula (MNI_xyz_: 40, -16, 6, *F*_(1,75)_ = 26.46, *p*_FWE_ < 0.01; Faces _Block_ _1_ > Faces _Block 3_), showing that high-lonely men exhibited increased insula habituation than high-lonely women. For the habituation to fearful faces a sex*loneliness interaction was observed in the right nucleus accumbens (MNI_xyz_: 18, 10, -12, *F*_(1,75)_ = 13.51, *p*_FWE_ = 0.01; Fearful _Block_ _1_ > Fearful _Block 3_) such that high-lonely women showed decreased habituation to fearful faces in contrast to high-lonely men. Furthermore, habituation to fearful faces compared to non-social stimuli (i.e., houses) was decreased in high-lonely woman compared to high-lonely men in the right nucleus accumbens (MNI_xyz_: 18, 8, -12, *F*_(1,75)_ = 9.91, *p*_FWE_ = 0.045; Fearful _Block 1 > Block 3_ > Houses _Block 1 > Block 3_). Additionally, left amygdala habituation (MNI_xyz_: -28, 0, -26, *F*_(1,75)_ = 15.69, *p*_FWE_ = 0.01; Faces _Block 1 > Block 3_ > Houses _Block 1 > Block 3_) to all faces relative to non-social stimuli was reduced in high-lonely women compared to high-lonely men. The opposite pattern was evident in low-lonely participants.

Furthermore, we observed a sex*loneliness interaction in functional connectivity with the right mOFC as seed region. In the habituation to social stimuli in contrast to non-social stimuli, high-lonely women showed higher coupling between the right mOFC and the right lateral occipital cortex (MNI_xyz_: 42, -42, 50, *k*_E(1,75)_ = 122, *p*_FWE_ < 0.01; Faces _Block 1 > Block 3_ > Houses _Block 1 > Block 3_) than high-lonely men. Furthermore, left amygdala connectivity with the left precentral gyrus (MNI_xyz_: -12, -32, 50, *k*_E(1,75)_ = 93, *p*_FWE_ = 0.02; Faces _Block 1 > Block 3_ > Houses _Block 1 > Block 3_) was decreased in high-lonely women in contrast to high-lonely men in the process of social stimuli habituation.

**Fear conditioning and extinction: reaction times**

A first RT analysis across conditioning and extinction showed that subjects reacted faster in the extinction in contrast to the conditioning phase (*F*_(1,64)_ = 104.30, *p* < 0.01, η_p_^2^ = 0.62). In addition, RTs to social stimuli were significantly faster than RTs to non-social stimuli (main effect of sociality type: *F*_(1,64)_ = 17.43, *p* < 0.01, η_p_^2^ = 0.21). No significant sex*loneliness interactions were observed (all *p*s > 0.05).

In the conditioning phase, subjects reacted faster in the second half than in the first half of conditioning (main effect of time: *F*_(1,64)_ = 29.88, *p* < 0.01, η_p_^2^ = 0.32). Furthermore, subjects showed significantly faster RTs to social stimuli than non-social stimuli (main effect of sociality: *F*_(1,64)_ = 34.47, *p* < 0.01, η_p_^2^ = 0.35). In addition a significant time*condition interaction was observed (*F*_(1,64)_ = 28.16, *p* < 0.01, η_p_^2^ = 0.31) showing that RTs in the CS- condition dropped faster than RTs in the CS+ condition, resembling the learning effect of conditioning.

A similar pattern emerged in the extinction phase with faster RTs in the second half of the experiment (main effect of time: *F*_(1,64)_ = 35.48, *p* < 0.01, η_p_^2^ = 0.36). Furthermore, we observed a significant time*condition interaction (*F*_(1,64)_ = 4.17, *p* = 0.045, η_p_^2^ = 0.06), whereby the decrease in RTs was more pronounced for the CS+ than for the CS-. In addition, high-lonely subjects exhibited higher RTs in the first half, but lower RTs in the second half of the extinction than low-lonely individuals (interaction effect time*loneliness: *F*_(1,64)_ = 6.75, *p* = 0.01, η_p_^2^ = 0.10). For RTs see **Tab.** **S4**.

**Fear conditioning and extinction: skin conductance response**

The analysis of SCRs during the COND/EXT fMRI paradigm revealed higher magnitudes in response to the CS+ compared to the CS- (*F*_(1,66)_ = 5.80, *p* = 0.02, η_p_^2^ = 0.08), as well as higher magnitudes across conditions in the conditioning phase than in the extinction phase (main effect of task: *F*_(1,66)_ = 4.01, *p* = 0.049, η_p_^2^ = 0.06). Neither sex nor loneliness significantly affected SCR magnitudes across sociality and time conditions (all *p*s > 0.05).

Furthermore, SCR magnitudes across conditions were significantly higher in the first eight trials of conditioning (*F*_(1,71)_ = 45.68, *p* < 0.01, η_p_^2^ = 0.39) and extinction (*F*_(1,67)_ = 8.20, *p* = 0.01, η_p_^2^ = 0.11) than in the last eight trials. In addition, high-lonely men showed increased SCR magnitudes to non-social stimuli during extinction in contrast to low-lonely men (interaction sociality * loneliness * sex: *F*_(1,67)_ = 6.45, *p* = 0.01, η_p_^2^ = 0.09).

**Fear conditioning and extinction: fMRI task effects**

Comparing the COND and EXT phases, we found higher activations in clusters involving the superior temporal gyrus and precentral gyrus (COND _CS+ > CS-_ > EXT _CS+ > CS-_, cf. **Tab.** **S5**) at the whole-brain level. Additional ROI analyses revealed higher activations in the amygdala (L: MNI_xyz_: -26, -4, -12, *t*_(75)_ = 4.98, *p*_FWE_ < 0.01; R: MNI_xyz_: 22, 0, -12, *t*_(75)_ = 4.42, *p*_FWE_ < 0.01), ACC (L: MNI_xyz_: 0, 16, 30, *t*_(75)_ = 6.85, *p*_FWE_ < 0.01; R: MNI_xyz_: 2, 18, 28, *t*_(75)_ = 7.05, *p*_FWE_ < 0.01), and insular cortex (L: MNI_xyz_: -36, 0, 10, *t*_(75)_ = 8.67, *p*_FWE_ < 0.01; R: MNI_xyz_: 34, -20, 18, *t*_(75)_ = 8.98, *p*_FWE_ < 0.01; COND _CS+ > CS-_ > EXT _CS+ > CS-_).

In the conditioning phase, ROI analyses confirmed significantly higher activations to the CS+ in the amygdala (L: MNI_xyz_: -18, -2, -12, *t*_(75)_ = 4.70, *p*_FWE_ < 0.01; R: MNI_xyz_: 20, 0, -12, *t*_(75)_ = 4.85, *p*_FWE_ < 0.01), ACC (L: MNI_xyz_: 0, 16, 28, *t*_(75)_ = 9.25, *p*_FWE_ < 0.01; R: MNI_xyz_: 2, 12, 28, *t*_(75)_ = 9.45, *p*_FWE_ < 0.01), and insula (L: MNI_xyz_: -28, 20, 10, *t*_(75)_ = 12.24, *p*_FWE_ < 0.01; R: MNI_xyz_: 36, 16, 4, *t*_(75)_ = 10.92, *p*_FWE_ < 0.01; COND _CS+ > CS-_), as well as decreased activations to the CS+ in the mOFC (L: MNI_xyz_: -6, 42, -14, *t*_(75)_ = 4.63, *p*_FWE_ < 0.01; R: MNI_xyz_: 12, 44, -8, *t*_(75)_ = 4.84, *p*_FWE_ < 0.01; COND _CS+ < CS-_). In the EXT phase, whole-brain analysis revealed that the CS+ induced activations in the right insula, supramarginal gyrus, superior frontal gyrus, and supplementary motor area (EXT _CS+ > CS-_; cf. **Tab. S5**). In addition, ROI analyses revealed higher insula (L: MNI_xyz_: -30, 18, 8, *t*_(75)_ = 4.11, *p*_FWE_ = 0.03; R: MNI_xyz_: 32, 18, -8, *t*_(75)_ = 5.75, *p*_FWE_ < 0.01;) and ACC (L: MNI_xyz_: 2, 28, 28, *t*_(75)_ = 4.48, *p*_FWE_ = 0.01; R: MNI_xyz_: 8, 22, 26, *t*_(75)_ = 5.34, *p*_FWE_ < 0.01; EXT _CS+ > CS-_) reactivity to the CS+.

**Fear conditioning and extinction: fMRI effects**

In addition, ROI analyses, we found a sex*loneliness interaction in the activity of the left mOFC to social fear stimuli in the early phase of conditioning compared to that of extinction (MNI_xyz_: -12, 44, -8, *F*_(1,72)_ = 19.89, *p*_FWE_ = 0.01; COND _CS+ social > CS- social_ > EXT _CS+ social > CS- social_) such that high-lonely men showed reduced mOFC responses compared to high-lonely women and the opposite pattern emerged in low-lonely individuals. The same effect in the mOFC (MNI_xyz_: -12, 42, -6, *F*_(1,72)_ = 15.51, *p*_FWE_ = 0.04; COND _CS+ social > CS- social_ > EXT _CS+ social > CS- social_) was evident across all trials.

In the first half of the conditioning phase, an additional sex*loneliness interaction was found for hippocampal responses to social stimuli. High-lonely men exhibited stronger hippocampus activity in response to social threat cues compared with high-lonely women (MNI_xyz_: 26, -42, 2, *F*_(1,72)_ = 21.36, *p*_FWE_ = 0.01; COND _CS+ social > CS+ non-social_ > COND _CS- social > CS- non-social_).

In addition, over all trials of the extinction phase a sex*loneliness interaction was observed in the right and left ACC in response to social CS+ compared to non-social CS+ (L: MNI_xyz_: 0, 34, 20, *F*_(1,72)_ = 22.37, *p*_FWE_ = 0.01; R: MNI_xyz_: 2, 32, 20, *F*_(1,72)_ = 18.94, *p*_FWE_ = 0.02; EXT _CS+_ _social > CS+ non-social_ > EXT _CS-_ _social > CS- non-social_) showing that high-lonely men exhibited decreased activity in contrast to high-lonely women. A sex*loneliness interaction was also observed for ACC responses to social CS+ relative to social CSs- (L: MNI_xyz_: 0, 34, 18, *F*_(1,72)_ = 21.48, *p*_FWE_ = 0.01; R: MNI_xyz_: 2, 32, 20, *F*_(1,72)_ = 17.34, *p*_FWE_ = 0.04; EXT _CS+ social > CS- social_).

Furthermore, functional connectivity analyses revealed another sex*loneliness interaction. In the extinction phase, high-lonely men showed stronger coupling between the left hippocampus as seed region and the left superior frontal gyrus (MNI_xyz_: -22, -4, 56, *k*_E(1,72)_ = 123, *p*_FWE_ = 0.01) compared to high-lonely women for social stimuli (EXT _CS+_ _social > CS+ non-social_ > EXT _CS-_ _social > CS- non-social_). In addition, for social stimuli relative to non-social stimuli over all trials stronger coupling between the right amygdala and the frontal cortex (MNI_xyz_: 30, 50, -20, *k*_E(1,72)_ = 120, *p*_FWE_ = 0.01) was evident in high-lonely men compared to high-lonely women (EXT _CS+_ _social > non-social_ > EXT _CS-_ _social > non-social_). Moreover, in the first half of conditioning and extinction trials, high-lonely men showed an increased coupling between the right insula as seed region and the right middle frontal gyrus (MNI_xyz_: 40, 14, 38, *k*_E(1,72)_ = 111, *p*_FWE_ = 0.01) in contrast to high-lonely women who exhibited decreased coupling (COND _CS+ social > CS- social_ > EXT _CS+ social > CS- social_).

**Brain behavior connections**

Parameter estimates of significant clusters derived from the full factorial models were correlated with the number of intrusions. Only right amygdala habituation to fearful faces in the emotional face-matching task correlated significantly with the number of intrusions (*r*_(76)_ = -0.22 *p* = 0.049; Fearful _Block_ _1_ > Fearful _Block 3_).

**Supplementary Tables**

**Table S1.** Hormonal blood concentrations at baseline

|  | **High-lonely** | | **Low-lonely** | |
| --- | --- | --- | --- | --- |
|  | **Men** | **Women** | **Men** | **Women** |
| Testosterone | 4.29 (1.50) | 0.66 (1.31) | 5.05 (1.44) | 0.25 (0.12) |
| Progesterone | 0.18 (0.13) | 3.41 (6.22) | 0.14 (0.05) | 0.31 (0.52) |
| Estradiol | 25.10 (8.33) | 109.57 (113.00) | 25.49 (5.32) | 33.53 (22.39) |
| DHEAS | 3.54 (1.12) | 2.81 (1.43) | 3.70 (1.17) | 2.45 (1.22) |
| LH | 5.17 (1.65) | 6.95 (6.41) | 5.32 (1.36) | 6.34 (2.32) |
| FSH | 4.18 (2.37) | 4.51 (1.98) | 3.11 (1.45) | 4.35 (2.45) |

*Notes.* The table shows means and standard deviations in brackets. DHEAS concentrations in µg/l. Progesterone and testosterone concentrations are shown in ng/ml. Estradiol concentrations are shown in pg/ml. LH are shown in U/l. FSH are shown in mlU/ml. Abbreviations: LH, luteinizing hormone, FSH, follicle-stimulating hormone, DHEAS, dehydroepiandrosterone

**Table** **S2**. Reaction times and correct responses in the emotional face-matching task across groups

|  | **RTs (s)** | | **CRs (%)** | |
| --- | --- | --- | --- | --- |
|  | **Mean** | **SD** | **Mean** | **SD** |
| Fearful | 1.23 | 0.23 | 98.72 | 3.73 |
| Happy | 1.32 | 0.28 | 98.63 | 3.29 |
| Neutral | 1.30 | 0.28 | 98.63 | 3.11 |
| House | 1.37 | 0.28 | 98.55 | 2.97 |

*Notes.* The mean reaction times (RTs) and standard deviations (SD) are shown in seconds. Correct response rates (CRs) are shown as percentages.

**Table S3.** Whole-brain findings of emotional face habituation across groups

| **Region** | **Right/left** | **Cluster size** | | **Peak *t*** | | **MNI coordinates** | | | |
| --- | --- | --- | --- | --- | --- | --- | --- | --- | --- |
|  |  |  |  |  |  | **x** | | **y** | **z** |
| **Habituation fearful faces** |  |  | |  | |  | |  |  |
| Superior frontal gyrus | right | 187 | | 5.41 | | 20 | | 12 | 58 |
| Supramarginal gyrus | right | 546 | | 5.38 | | 56 | | -42 | 28 |
| **Habituation all faces** |  |  | |  | |  | |  |  |
| Cuneus | left | 6978 | | 6.11 | | 0 | | -86 | 30 |
| **Habituation all faces > habituation house** | | |  | |  | |  |  |  |
| Lingual gyrus | left | 1607 | | 6.14 | | -20 | | -56 | -12 |
| Cuneus | left | 1129 | | 6.09 | | -6 | | -92 | 28 |
| Fusiform gyrus | right | 1621 | | 5.64 | | 24 | | -50 | -14 |

*Notes.* An initial cluster-forming height threshold of *p* < 0.001 was used. Only cluster with FWE-corrected *p*s < 0.05 at the peak level are listed. Abbreviations: MNI, Montreal Neurological Institute

**Table S4.** Reaction times and contingency ratings for the conditioning and extinction tasks across groups

|  | **RTs (s)** | | **Contingency rating** | |
| --- | --- | --- | --- | --- |
|  | **Mean** | **SD** | **Mean** | **SD** |
| **COND first half** |  |  |  |  |
| CS+ Social | 1.35 | 0.43 | 0.44 | 0.59 |
| CS+ Non-Social | 1.45 | 0.35 | 0.26 | 0.56 |
| CS- Social | 1.37 | 0.37 | -0.63 | 0.46 |
| CS- Non-Social | 1.54 | 0.49 | -0.57 | 0.52 |
| **COND second half** |  |  |  |  |
| CS+ Social | 1.23 | 0.43 | 0.70 | 0.52 |
| CS+ Non-Social | 1.40 | 0.43 | 0.56 | 0.55 |
| CS- Social | 1.16 | 0.30 | -0.88 | 0.32 |
| CS- Non-Social | 1.32 | 0.43 | -0.74 | 0.51 |
| **EXT first half** |  |  |  |  |
| CS+ Social | 1.11 | 0.48 | -0.50 | 0.54 |
| CS+ Non-Social | 1.11 | 0.37 | -0.54 | 0.52 |
| CS- Social | 1.08 | 0.40 | -0.87 | 0.29 |
| CS- Non-Social | 1.09 | 0.42 | -0.90 | 0.35 |
| **EXT second half** |  |  |  |  |
| CS+ Social | 1.00 | 0.53 | -0.92 | 0.31 |
| CS+ Non-Social | 0.99 | 0.46 | -0.95 | 0.26 |
| CS- Social | 0.95 | 0.38 | -0.94 | 0.32 |
| CS- Non-Social | 1.00 | 0.48 | -0.98 | 0.10 |

*Notes.* The mean reaction times (RTs) and standard deviations (SD) are shown in seconds. Contingency ratings vary between -1 and +1, with -1 indicating CS- and +1 indicating CS+. Abbreviations: COND, conditioning; CS+, fear-associated conditioned stimulus; CS-, non-fear-associated conditioned stimulus; EXT, extinction; first half, first eight trials of the task; second half, second eight trials of the task.

**Table S5.** Whole-brain findings for the fear condition and extinction tasks across groups

| **Region** | **Right/left** | **Cluster size** | **Peak *t*** | **MNI coordinates** | | |
| --- | --- | --- | --- | --- | --- | --- |
|  |  |  |  | **x** | **y** | **z** |
| **COND _CS+ > CS-_ ^1^** |  |  |  |  |  |  |
| Postcentral gyrus | left | 34197 | 13.34 | -58 | -22 | 26 |
| Precuneus | right | 1016 | 7.40 | 14 | -66 | 38 |
| **EXT _CS+ > CS-_ ^2^** |  |  |  |  |  |  |
| Insula | right | 660 | 5.75 | 32 | 18 | -8 |
| Supramarginal gyrus | right | 150 | 5.53 | 58 | -42 | 26 |
| Supplementary motor area | right | 217 | 5.47 | 10 | 14 | 56 |
| Superior frontal gyrus | right | 539 | 5.34 | 8 | 22 | 26 |
| **COND _CS+ > CS-_ > EXT _CS+ > CS-_ ^3^** | |  |  |  |  |  |
| Superior temporal gyrus | left | 23920 | 12.29 | -46 | -34 | 22 |
| Precentral gyrus | left | 579 | 6.53 | -44 | -6 | 52 |
| **EXT _CS+ > CS-_ > COND _CS+ > CS-_** | | | | |  |  |
| No significant effects |  |  |  |  |  |  |

*Notes.* An initial cluster-forming height threshold of *p* < 0.001 was used. Only cluster with FWE-corrected *p*s < 0.05 on peak level are listed. ^1^ ROI analyses revealed increased activations in the amygdala, anterior cingulate cortex and insula, as well as decreased activation in the medial prefrontal cortex. ^2^ ROI analyses revealed increased activations in the anterior cingulate and insula cortex. ^3^ ROI analyses revealed increased activations in the insula and anterior cingulate cortex, as well as in the amygdala. Abbreviations: MNI, Montreal Neurological Institute; COND, conditioning; CS+, fear-associated conditioned stimulus; CS-, non-fear-associated conditioned stimulus; EXT, extinction.

**Table S6.** Baseline differences between groups

|  | **High-lonely** | | **Low-lonely** | |
| --- | --- | --- | --- | --- |
|  | **Women**  **(n = 20)** | **Men**  **(n = 27)** | **Women**  **(n = 18)** | **Men**  **(n = 17)** |
| Loneliness ^a)^ | 54.60 (5.62) | 55.19 (3.53) | 23.56 (1.20) | 24.06 (1.03) |
| Depressive symptoms ^b)^ | 4.25 (3.51) | 3.85 (3.91) | 2.11 (3.64) | 1.53 (2.15) |
| Social anxiety ^c)^ | 22.20 (17.20) | 22.52 (18.99) | 13.39 (9.85) | 11.82 (15.40) |
| Childhood maltreatment ^d)^ | 35.00 (9.43) | 38.44 (10.06) | 32.11 (15.32) | 29.47 (5.30) |
| Alexithymia ^e)^ | 41.15 (9.53) | 46.22 (10.43) | 32.39 (6.46) | 34.29 (6.54) |
| Social support ^f)^ | 60.40 (9.50) | 52.11 (12.88) | 68.11 (3.10) | 65.59 (12.88) |
| Perceived stress ^g)^ | 13.25 (7.09) | 12.96 (6.48) | 8.78 (5.11) | 7.35 (4.64) |
| Trait anxiety ^h)^ | 36.95 (7.71) | 40.15 (9.82) | 27.67 (5.13) | 26.35 (4.76) |
| Social network ^i)^ |  |  |  |  |
| Numbers | 18.35 (9.18) | 14.04 (5.40) | 21.22 (7.58) | 19.35 (7.31) |
| Roles | 5.30 (1.56) | 4.56 (1.05) | 5.78 (1.44) | 5.65 (1.62) |
| Networks | 1.80 (1.40) | 1.33 (1.00) | 2.22 (1.06) | 2.06 (1.20) |

*Notes.* Values are the mean and SD. ^a)^ Participants were pre-stratified and assigned to the high- or low-lonely group using the UCLA Loneliness Scale (UCLA-L). High-lonely participants had a score equal or above 50 while low-lonely participants had a score equal or below 25; ^b)^ Depressive symptoms were measured with the Beck Depression Inventory, Version II (BDI); ^c)^ Social anxiety was assessed with the Liebowitz Social Anxiety Scale (LSAS); ^d)^ Childhood traumata was measured using the Childhood Trauma Questionnaire (CTQ); ^e)^ Alexithymic symptoms were assessed by the Toronto Alexithymia Scale (TAS); ^f)^ Social Support was measured with the Social Support Questionnaire ([Fragebogen zur sozialen Unterstützung] ;F-SozU); ^g)^ Perceived stress was quantified by the perceived stress scale (PSS-10); ^h)^ Trait anxiety was assessed by the State Trait Anxiety Inventory (STAI); ^i)^ Social network was characterized using the Social Network Index to assess the number of diverse social roles, networks, and the total number of people the participants talked to regularly.

**Table S7.** Whole-brain task effects of the emotional face-matching task across groups

| **Region** | **Right/left** | **Cluster size** | **Peak *t*** | **MNI coordinates** | | |
| --- | --- | --- | --- | --- | --- | --- |
|  |  |  |  | **x** | **y** | **z** |
| **Faces > Houses** |  |  |  |  |  |  |
| Calcarine sulcus | right | 2878 | 11.28 | 24 | -96 | 0 |
| Hippocampus | right | 1407 | 10.10 | 20 | -6 | -12 |
| Amygdala | left | 919 | 9.20 | -20 | -6 | -14 |
| Fusiform gyrus | left | 680 | 8.22 | -40 | -50 | -18 |
| Medial orbital frontal gyrus | left | 535 | 6.39 | -2 | 40 | -12 |
| Inferior occipital lobe | left | 760 | 6.35 | -52 | -64 | -16 |
| Precuneus | right | 1682 | 5.78 | 4 | -56 | 28 |
| Inferior frontal gyrus (triangularis) | right | 363 | 5.30 | 44 | 16 | 24 |
| **Fearful > Houses** |  |  |  |  |  |  |
| Middle temporal gyrus | right | 2965 | 10.67 | 52 | -62 | 8 |
| Fusiform gyrus | left | 705 | 8.83 | -40 | -52 | -18 |
| Amygdala | left | 732 | 8.50 | -20 | -6 | -14 |
| Thalamus | right | 1335 | 7.70 | 20 | -6 | 12 |
| Medial orbital frontal gyrus | left | 1259 | 6.78 | -2 | 40 | -12 |
| Middle temporal gyrus | left | 1177 | 6.69 | -54 | -64 | 14 |
| Precuneus | right | 2791 | 5.93 | 4 | -60 | 30 |
| Inferior frontal gyrus (triangularis) | right | 276 | 5.43 | 40 | 18 | 22 |
| Inferior frontal gyrus (triangularis) | left | 304 | 5.42 | -42 | 14 | 28 |
| Precentral gyrus | right | 131 | 5.27 | 52 | 0 | 48 |
| Middle temporal gyrus | left | 260 | 5.24 | -50 | -14 | -14 |
| Cerebellum | left | 91 | 5.22 | -10 | -82 | -36 |
| **Fearful > Neutral** |  |  |  |  |  |  |
| Middle temporal gyrus | left | 654 | 5.51 | -50 | -48 | 10 |
| *Notes.* An initial cluster-forming height threshold of *p* < 0.001 was used. Only cluster with FWE-corrected *p*s < 0.05 on peak level are listed. Abbreviations: MNI, Montreal Neurological Institute | | | | |  |  |

**Supplementary Figures**

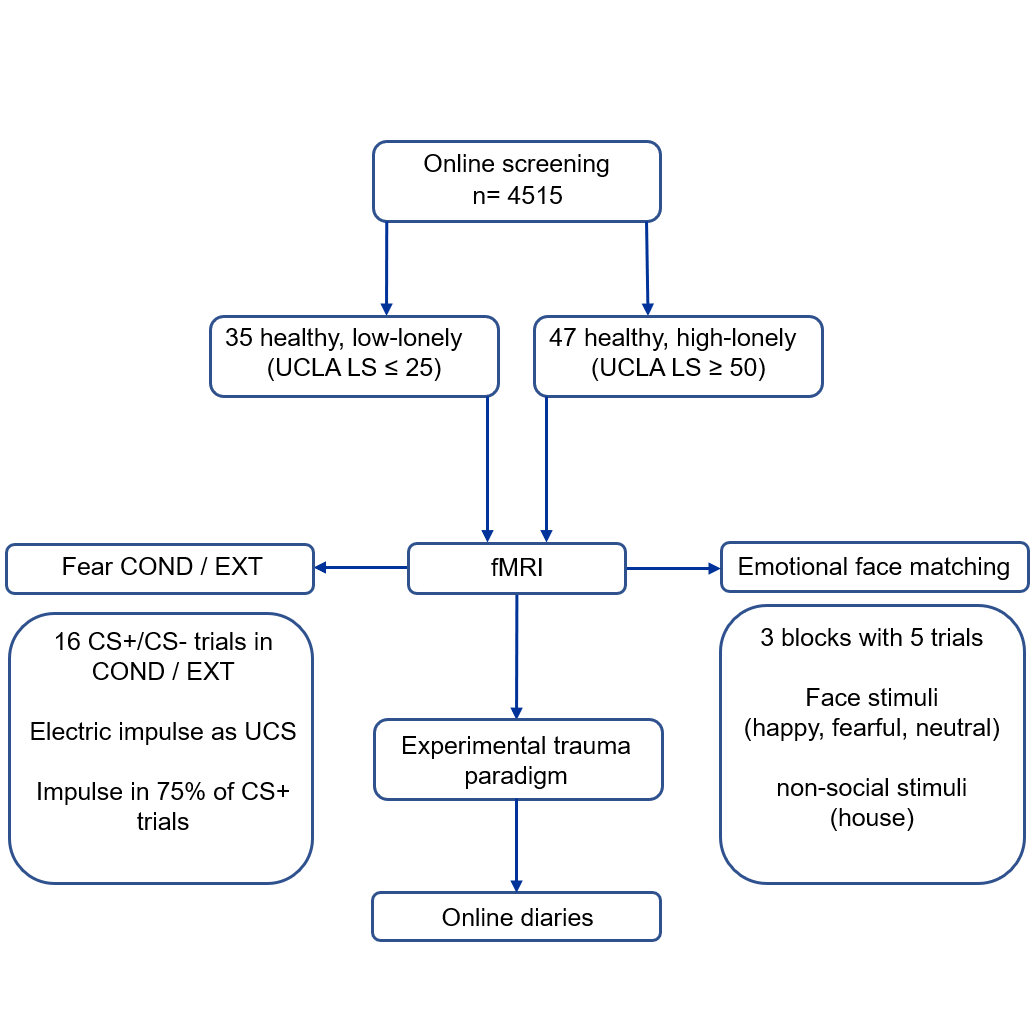

**Fig. S1.** Schematic overview of the study protocol. Subjects were recruited via an online questionnaire (n = 4515). Ninety-seven participants were invited for screening. In the screening session, the medical history and questionnaire data were assessed. Fifteen participants had to be excluded after the screening session because they were not eligible for enrollment, resulting in a final sample of 82 healthy subjects (38 women, mean age ± standard deviation [SD]: 26.39 ± 5.83 years; high-lonely: n = 47; low-lonely: n = 35). The testing session consisted of an fMRI scan containing a high-resolution structural scan, a fear conditioning (COND) / extinction (EXT) paradigm, and an emotional face matching paradigm. Following the fMRI scan, subjects viewed a trauma video. To measure intrusive thoughts, subjects completed online diaries in the three days following the video session. Abbreviations: COND, conditioning; CS+, fear-associated conditioned stimulus; CS-, non-fear-associated conditioned stimulus; EXT, extinction; UCLA LS, UCLA loneliness scale; UCS, unconditioned stimulus.
